# Pre-analytical delay as a dominant confounder in blood RNA-seq: Rapid ex vivo gene expression changes in EDTA blood

**DOI:** 10.64898/2026.06.27.734945

**Authors:** Kalle Günther, Ioanna Andreou, Daniel Kim, Jonathan M. Shaffer, Markus Sprenger-Haussels

## Abstract

Although the impact of delayed processing on gene expression in EDTA blood has been well documented using targeted assays and microarray platforms, the emergence of next-generation RNA sequencing (RNA-seq) has not yet been leveraged to systematically compare these effects against stabilized whole blood collection systems. Notably, no study has performed a time course RNA-seq analysis with human bulk RNA of matched EDTA and PAXgene blood RNA samples drawn from the same subjects. EDTA is still widely used for gene expression analysis studies. Yet, the genome-wide dynamics by which EDTA blood transcriptomes deviate from a stabilized reference over time remain poorly defined. This represents an important methodological gap, given the increasing reliance on RNA-seq for biomarker discovery, clinical transcriptomics and diagnostics.

## Introduction

Peripheral blood is a revealing source of transcriptional information to assess unique gene expression profiles potentially associated with disease predictions, detection or monitoring [1]. It is used in many basic scientific research studies of human diseases, in pharmacogenomic studies and clinical trials [1]. To ensure reliable and accurate results in those studies, it is of utmost importance to preserve human whole blood specimens and their transcriptional profiles after blood draw.

Gene expression patterns and levels differ between cells depending on a cell’s actual state [1]. Many factors can influence RNA quality, change gene expression or even lead to RNA degradation affecting downstream analysis. These include the blood collection method, presence or absence of anticoagulants, stabilization technology, sample handling, storage, temperature upon sample processing and the processing methods themselves. It is therefore important to evaluate how blood collection, storage and extraction methodologies influence gene expression profiling as analyzed in downstream processes such as RT-qPCR and RNA sequencing (RNA-seq). It is possible that bias may result from the confounding effects of the above-mentioned workflow steps, especially when considering a donor’s individual genetic constitution or changes with respect to a disease [1,2]. Any subtle changes in gene expression readout potentially caused by a pre-analytical step may lead to false changes in the diagnostic, prognostic and predictive markers used to analyze a disease [1].

There is evidence that conventional dipotassium ethylenediaminetetraacetic acid (K_2_ EDTA) blood collection tubes do not accurately preserve gene expression levels in blood samples after sample collection [3–5], but the transcriptome-wide extent of the change remains unknown. The PAXgene Blood RNA Tube (IVD; PreAnalytiX, Hombrechtikon, Switzerland; hereafter referred to as PAXgene tube) is a single tube for collection, storage and transport of whole blood and immediate stabilization of intracellular RNA at collection site [6].

Additives in the PAXgene Blood RNA Tube effectively trigger the lysis of blood cells and subsequently protect RNA molecules in the sample from degradation by RNases and other enzymes. This minimizes ex vivo changes in gene expression due to gene induction and downregulation. The result is immediate and allows stabilization of RNA for up to 3 days at room temperature, up to 5 days at 2–8°C and long-term stabilization of up to 11 years at −20°C or −70°C [7].

RNA-seq is a continuously evolving area of transcriptomics. It enables researchers to quantify gene expression, discover novel transcripts and analyze alternative splicing events that enable a broad variety of research, translational and clinical applications [8–11]. Data from RNA-seq enables the interpretation of gene expression changes associated with cellular function, mechanistic study of biological pathways, biomarker discovery, characterization of druggable pathways and disease diagnostics [9,12–14]. Inherently, gene expression is a dynamic process that can be rapidly altered by the ex vivo conditions associated with specimen collection, transport, storage, archiving and processing. Stressors can induce artifactual changes in transcript abundance, which confound the interpretation of gene expression data [15,16] and render research and diagnostic findings unreliable [16]. Immediate specimen stabilization therefore is essential not only to preserve RNA integrity but also to maintain the transcriptome’s in vivo state.

Degradation can begin within minutes of blood lysis if proper precautions are not taken, which can negatively impact RNA-seq library preparation and data quality [17]. It can cause biased read coverage – particularly favoring the 3’ ends of transcripts – and reduce the diversity and complexity of the sequencing library [18]. Collectively, RNA-seq’s accuracy and reproducibility are intimately associated with the quality of the input RNA. Thus, a prerequisite for accurate and reproducible RNA-seq is that the specimen’s gene expression profile does not change during storage or transport; any changes here would distort the true, original expression patterns. While clinical laboratories are often focused on optimizing downstream assays, pre-analytical steps (sample collection, stabilization, transport and preparation) are of equal or greater importance [19,20].

In this study, we present a transcriptome-wide comparison of two widely used pre-analytical workflows for blood RNA analysis (using K_2_ EDTA tubes and the PAXgene Blood RNA Tube). Furthermore, we compare RNA sequencing results to relative quantification results obtained for individual, selected transcripts by RT-qPCR. We demonstrate that obtaining high-quality RT-qPCR and RNA-seq data begins at specimen collection. Immediate stabilization of RNA not only preserves molecular integrity but also ensures that the transcriptome reflects the true biological state at sample collection time, generating reproducible, biologically meaningful RNA-seq and RT-qPCR results.

## Materials and methods

The pre-analytical phase of the study adhered to ISO Standard 20186-1 [21].

### Study design

Blood was collected from nine consented healthy adults into multiple K_2_ EDTA Tubes (3 mL blood draw; BD, Franklin Lakes, NJ) and PAXgene Blood RNA Tubes (2.5 mL blood draw; PreAnalytiX, Hombrechtikon, Switzerland) per subject. One EDTA tube was processed immediately (t0). All other EDTA and PAXgene Blood RNA Tubes were stored at room temperature and processed at multiple time intervals up to 72 hours (3 days). Note: No PAXgene Blood RNA Tube was processed at t0 due to the minimum time of 2 hours required for sufficient RNA precipitation, while blood lysis occurs immediately upon contact between blood and the RNA stabilization solution in the tube. White blood cells (WBCs) were counted from the first EDTA tube of each subject without storage. At each timepoint, one PAXgene tube per subject was frozen at −20°C and one EDTA tube was used for total RNA isolation by combining acid guanidinium thiocyanate-phenol-chloroform based extraction according to Chomczynski and Sacchi [22] with further RNA purification by silica membrane-based cleanup. At the end of the time course, frozen PAXgene Blood RNA Tubes were thawed and subjected to RNA isolation with the PAXgene Blood RNA Kit (PreAnalytiX, Hombrechtikon, Switzerland) using both the manual protocol and the automated protocol on the QIAcube instrument (QIAGEN, Hilden, Germany) and also with the QIAsymphony PAXgene Blood RNA Kit on the QIAsymphony instrument (QIAGEN, Hilden, Germany) (three subjects per protocol). After processing of all specimens and RNA isolation, RNA samples were frozen at −20°C until analysis. RNA was quantified and assessed for quality by UV spectrophotometry and RNA integrity by miniaturized gel electrophoresis with RIN calculation. Gene expression was analyzed with real-time RT-qPCR of *FOS*, *IL1B*, *CXCL8* and *TP53* and with RNA sequencing. An overview of the time course experiment is shown in Fig 1.

**Fig 1.**
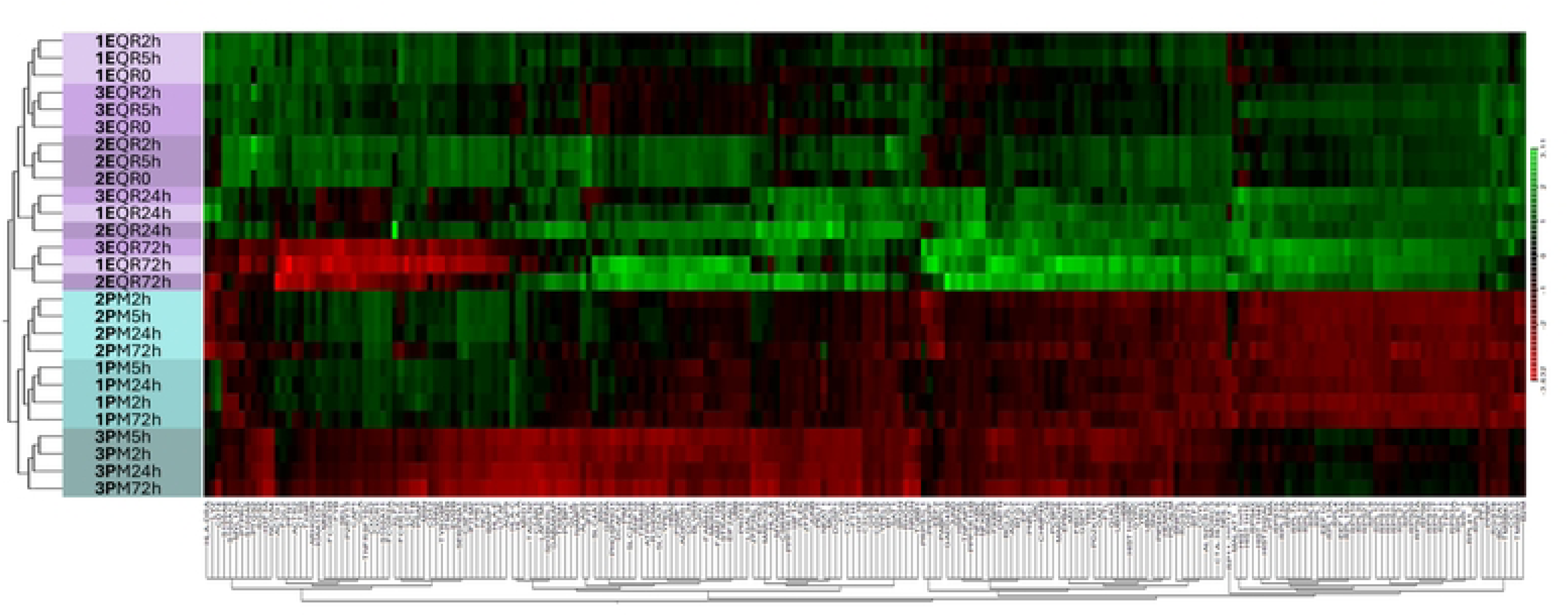
Flowchart of the RNA time course experiment setup. (A) Per subject, six EDTA tubes (E) and five PAXgene Blood RNA Tubes (P) were collected. EDTA tubes were either processed immediately [E*t_0_*] or stored for 2 to 72 hours (3 days) at room temperature [E*t_2 h_*-E*t_72 h_*]. (B) At the indicated test timepoints, specimens in PAXgene Blood RNA Tubes (P*t_2 h_*-P*t_72 h_*) were transferred to −20°C. (C) Specimens in EDTA tubes (E*t_0_*-E*t_72 h_*) were processed without freezing, while specimens in PAXgene Blood RNA Tubes were frozen and thawed before processing. Samples were subjected to RNA isolation with one of four protocols. (D) Resulting RNA was stored at −20°C. RNA isolation protocols: EQR: EDTA-QIAzol-RNeasy; PM: PAXgene Blood RNA Kit and manual processing; PQc: PAXgene Blood RNA Kit and QIAcube; PQs: QIAsymphony PAXgene Blood RNA Kit and QIAsymphony.

### Human blood specimens

In accordance with ethics approval granted by the ethics committee of North Rhine Westphalia, Germany (2007389), blood was collected via the company internal blood collection service in Hilden (Germany) under the supervision of the company’s medical officer on November 5, 2019. All subjects were volunteers and employees at QIAGEN’s site in Hilden (Germany) and were under constant medical review as stipulated by German employment law. The blood collection service accepted subjects aged between 18 and 68 years who underwent an additional, blood-donation specific medical examination before their first donation. The hematocrit value of all subjects was monitored monthly. Exclusion criteria for subjects were a hemoglobin value below 12.5 g/dL for women and 13.5 g/dL for men, a body weight below 50 kg, a body mass index ≤18, or a known pregnancy or illness. Prior to blood donation, the subjects gave their written consent. In accordance with the ethics committee approval, all blood specimens were anonymized before being handed over to the laboratory researchers.

Peripheral whole blood was collected by venipuncture and replicate specimens were drawn into six K_2_ EDTA Tubes and five PAXgene Blood RNA Tubes per subject. All blood draws were carried out according to the tube manufacturer’s instructions for use including 8–10 tube inversions after collection. Tubes were transported to the laboratory immediately without delay and processed directly.

WBCs were counted from EDTA blood samples using a DxH 500 Hematology Analyzer (Beckman Coulter, Brea, CA).

### RNA isolation methods

Total RNA was extracted from EDTA tubes either immediately or after storage at room temperature for different periods of time. Blood aliquots of 2 mL were mixed with QIAzol (QIAGEN, Hilden, Germany) and RNA was isolated according to the manufacturer’s instructions, with the following exceptions. RNA was not precipitated but purified by binding to a silica membrane. In detail, 0.5 volumes of ethanol (abs.) was added to the aqueous phase. The mixture was transferred to RNA binding spin columns of the RNeasy Mini Kit (QIAGEN) and the RNA cleanup protocol in the kit handbook was followed, including on-column DNase digestion steps. RNA was eluted twice with 40 µL RNase-free water (80 µL eluate volume).

For RNA isolation from specimens in PAXgene Blood RNA Tubes, tubes stored at −20°C were thawed for 2 hours at room temperature and specimens from three subjects each were subjected to RNA extraction using the manual protocol (PM) and the QIAcube automated protocol of the PAXgene Blood RNA Kit (PQc) and using the QIAsymphony PAXgene Blood RNA Kit according to the manufacturer’s instructions for use.

### RNA analysis methods

#### RNA quality and quantity

RNA was quantified and purity analyzed from RNA aliquots diluted in 10 mM Tris-HCl, (pH 7.5) by UV spectrophotometric measurements at 260, 280 and 320 nm with a SpectraMax Plus 384 instrument, which was correctly zeroed using the corresponding RNA elution solution, and SoftMax Pro Software V.3.1.2 (Molecular Devices, San Jose, CA). Absorption at 320 nm was used to subtract background absorption at 260 and 280 nm. RNA purity was calculated as absorbance ratio UV260/280 and RNA total yield based on UV260, dilution factor, extinction coefficient of RNA in buffered aqueous solutions and total elution volume. RNA integrity was analyzed by microcapillary gel electrophoretic RNA fragment length analysis using Eukaryote Total RNA NanoChips on the 2100 Bioanalyzer Instrument and RIN value calculation with Analysis Software 2100 Expert B.02.08.SI648 (Agilent Technologies, Santa Clara, CA) according to the manufacturer’s instructions.

#### Transcript level analyses by RT-qPCR

Relative differences of *FOS* and *IL1B* gene transcript levels between RNA samples were measured by real-time duplex reverse transcription quantitative PCR (RT-qPCR), normalized against 18S rRNA to compensate for pipetting inaccuracies. In addition, RT-qPCR assays targeting *CXCL8* and *TP53* gene transcripts were utilized as monoplex RT-qPCRs without normalization against 18S rRNA. RT-qPCR was performed with target specific primers and probes, 9 ng template RNA, each reaction, with 40 cycles on a Rotor-Gene Q cycler (QIAGEN) using the QuantiNova Probe RT-PCR Kit (QIAGEN), according to the kit handbook.

Performance of all assays was optimized, and *FOS* and *IL1B* assays were validated according to USP and CLSI guidelines to establish the method’s linear range, quantitation limit, specificity, selectivity and precision.

For each subject and RNA isolation protocol used, samples of the shortest blood storage time were used as reference samples to which all other samples of the set were compared, E*t_0_* for EDTA samples and P*t_2 h_* for PAXgene samples, as the stabilizer reagent of the PAXgene Blood RNA Tubes immediately lyses the cells and precipitation of RNA takes a minimum of 2 hours at room temperature before RNA isolation can be started (see PAXgene Blood RNA Kit Instructions For Use) [6].

Differences of gene expression levels between test and reference samples at a given test timepoint (T x) were calculated as ΔCt_T x_ for monoplex assays of *CXCL8* and *TP53* and as ΔΔCt_T x_ for duplex assays of *FOS* and *IL1B* according to:

ΔCt_T x_ = Ct_T ref_ – Ct_T x_ (monoplex assays) and
ΔΔCt_T x_ = ΔCt_T ref*_ – ΔCt_T x*_ (duplex assays), with
ΔCt_T ref*_ = Ct_T ref, 18S_ – Ct_T ref, target_
ΔCt_T x*_ = Ct_T x, 18S_ – Ct_T x, target_
18S: 18S rRNA sequence
target: *FOS*, *IL1B*, *CXCL8* or *TP53* gene transcript sequence
Ct: Cycle threshold of real time PCR reaction
ΔCt: Difference between two Ct values
ΔΔCt: Difference between two ΔCt values
T ref: Test timepoint serving as reference (*t_0_* for EDTA, *t_2 h_* for PAXgene)
T x: Test timepoint after x hours of blood specimen storage at room temperature

Using this approach, positive ΔCt_T x_ and ΔΔCt_T x_ values indicate gains and negative values indicate losses of transcripts. In accordance with the rule that a Ct difference of 1 means a doubling or halving of the transcript amount, depending on the mathematical sign, the data were used to calculate the x-fold changes compared to the initial level at the test time of the reference sample (set to 1) with: x-fold change _T x_ = 2^ΔCt,T x^ (for monoplex assays) and 2^ΔΔCt,T x^ (for duplex assays).

#### RNA sequencing

For each sample, RNA-seq libraries were constructed from 100 ng of total RNA utilizing the QIAseq Stranded RNA Library Kit (QIAGEN). Ribosomal RNA species (5S, 5.8S, 18S, 28S, 12S, 16S) were depleted using QIAseq FastSelect -rRNA HMR (QIAGEN), while globin mRNAs (HBA1, HBA2, HBB, HBD) were depleted with QIAseq FastSelect -Globin HMR (QIAGEN). In brief, library preparation involved fragmentation and targeted depletion of ribosomal RNA and globin transcripts, followed by first-strand cDNA synthesis. Second-strand synthesis, end-repair and A-tailing were conducted concurrently, followed by a magnetic bead purification step.

Strand-specific adapter ligation was then performed, again followed by bead cleanup. Library amplification and unique dual indexing (UDIs) for each sample were achieved via high-fidelity PCR, after which a final bead purification was conducted. Quality control of the resulting libraries was assessed using the TapeStation system with the D1000 ScreenTape Assay (Agilent Technologies, Santa Clara, CA, USA) to verify appropriate fragment size distribution, absence of adapter dimers and overall suitability for next-generation sequencing (NGS).

Twenty-seven libraries were multiplexed in each NextSeq 550 (Illumina, Inc., San Diego, CA, USA) run, with three sequencing runs performed in total; each sample received an average depth exceeding 25 million reads. Sequencing data were subsequently mapped and analyzed using the QIAGEN CLC Genomics Workbench (QIAGEN, Hilden, Germany). Pearson correlation was used to assess method consistency. Gene expression data were analyzed using PCA, volcano plots, and heat maps (RPKM, hierarchical clustering) to assess global patterns, differential expression, and sample relationships across stabilization conditions, time points, and donors. Differential expression analysis used outlier down-weighting with thresholds of adjusted *p* < 0.05 and | log₂ fold change | ≥ 1. Mitochondrial reads were excluded to prevent bias from high and variable mitochondrial transcript abundance [23].

### Statistical analyses

Data analysis was performed using GraphPad Prism V8.4.3 (GraphPad Software, LLC) for descriptive statistics and statistical tests, unless otherwise specified. Results were presented as mean ± SD, median with percentiles and single data points outside the given percentiles as described in the figures.

For each group of RNA samples isolated from EDTA (E) and PAXgene Blood RNA Tubes (P), or subgroup per RNA isolation protocol (EQR, PM, PQc, PQs) and RT-qPCR data (ΔCt_T x_, ΔΔCt_T x_) per test timepoint, data were tested for normal distribution by applying the Kolmogorov-Smirnov test with a *p*-level of 0.05.

For normally distributed data sets, one-way ANOVA was used to compare groups. Other sets were analyzed using nonparametric tests, with the Mann-Whitney U test for comparison of two groups and the Kruskal-Wallis H test for comparison of more than two groups.

Analyses were performed to identify significant differences between groups, considering a *p*-value of < 0.05 as statistically significant. Levels of significance are given as \**p* < 0.05, \*\**p* < 0.01, \*\*\**p* < 0.001, \*\*\*\**p* < 0.0001 and *p* ≥ 0.05 (*p*=*ns,* not significant).

Statistically significant differences of *FOS* and *IL1B* transcript levels (gains and losses of transcripts) between individual samples of the same subject and workflow (same blood tube and RNA isolation protocol) as measured by RT-qPCR are indicated by ΔΔCt_T x_ values greater than three times the method’s precision established by method validation (3 sigma; *p* < 0.0027).

## Results

### Assessment of RNA quantity and quality

A total of 99 blood specimens were collected from nine subjects. The time between blood collection and the start of RNA isolation from all EDTA samples without storage (E*_t0_* reference time point, transport time only between doctor’s office and laboratory) was on average 7.4 ± 2.5 minutes (mean and 1 SD), ranging from 4 to 12 minutes. WBC counts were determined from one EDTA tube per subject and the average number of WBC was 6.1 ± 1.3 x 10^6^ (mean ± SD), ranging from 4.6 x 10^6^ to 8.7 x 10^6^ WBC per mL of blood.

RNA yield ranged from 5.1 to 18.0 µg RNA for EDTA and from 4.3 to 17.3 µg RNA for PAXgene blood, while the median total yield of all EDTA and PAXgene samples was 8.4 µg and 7.4 µg, respectively (Fig 2).

**Fig 2.**
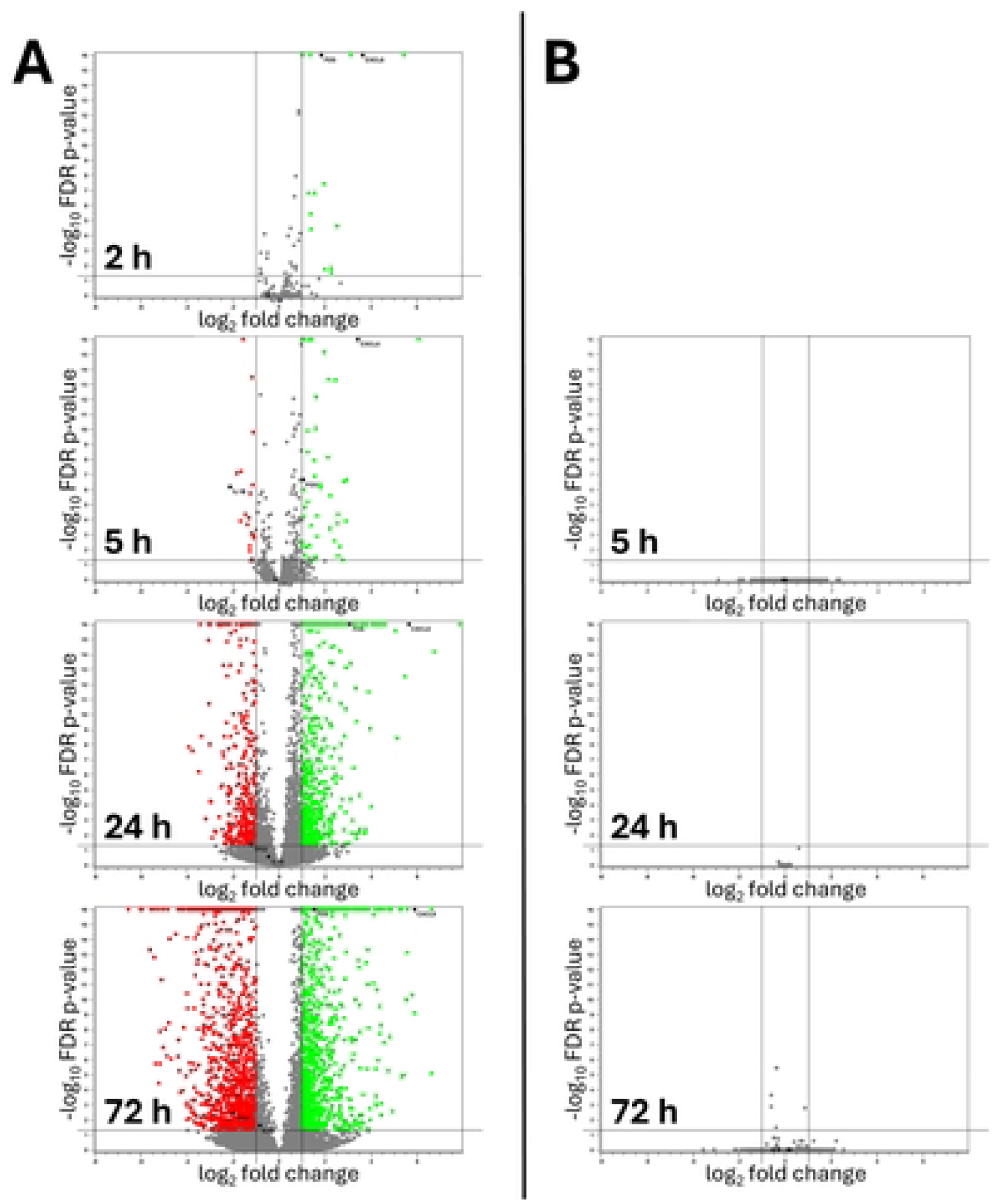
RNA quantity control: Absolute total RNA yields. RNA yield of all samples from all subjects is shown as individual data points and median per blood tube used for RNA isolation. E: EDTA tube specimens processed with EQR protocol (n = 54); P: PAXgene Blood RNA Tube specimens processed with PM, PQc and PQs protocol (n = 45). Statistical significance of difference between groups is indicated as \**p* < 0.05.

The median relative yield of PAXgene samples normalized to matching EDTA samples of the same subject and identical blood storage duration prior to RNA isolation was 92% for samples from all three RNA isolation protocols with PAXgene Blood RNA Tubes combined and 90%, 93% and 94% for PM, PQc and PQs, with no significant difference between the isolation protocols (Fig 3).

**Fig 3.**
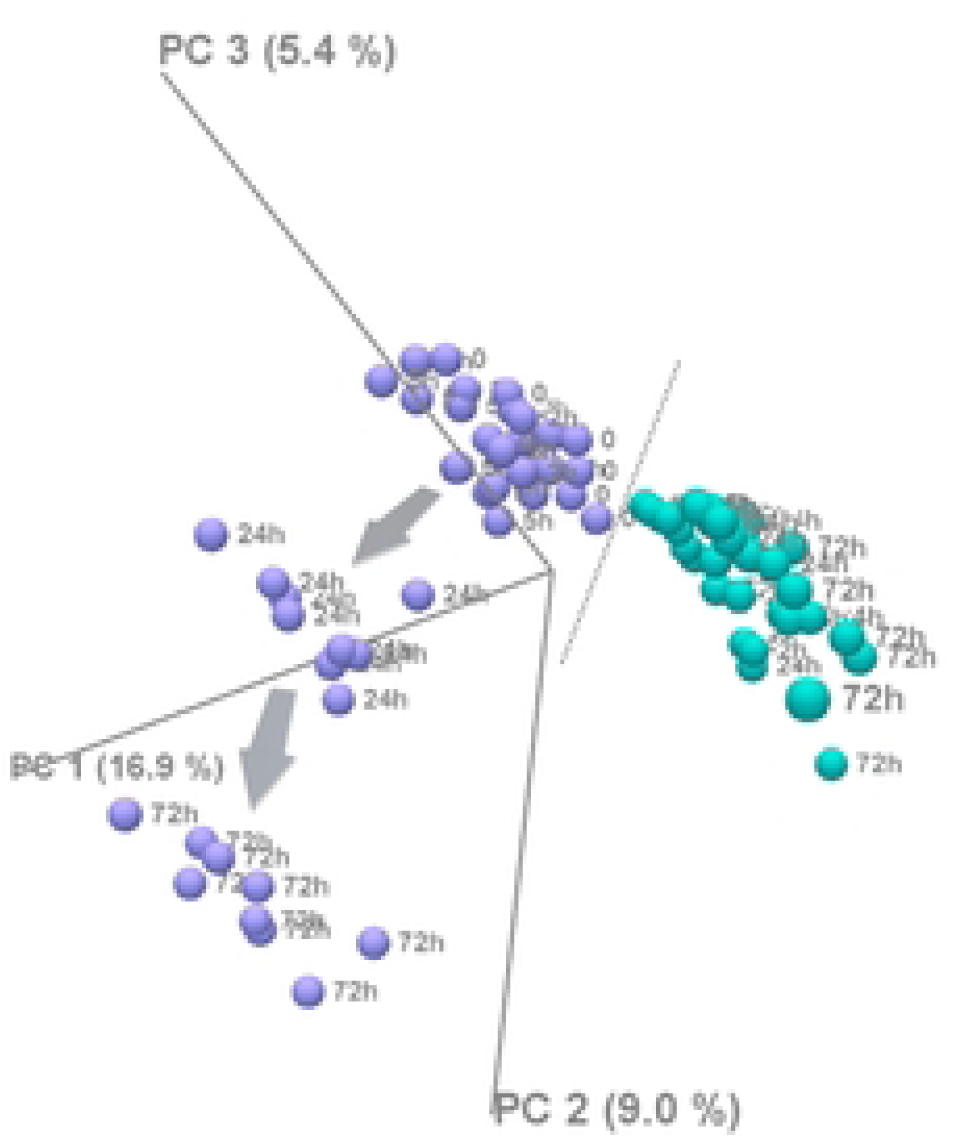
RNA quantity control: Relative total RNA yields of PAXgene samples normalized to matching EDTA samples. Relative RNA yield of each PAXgene sample was calculated based on the EDTA sample matching the same subject and blood storage test timepoint. Distribution of relative RNA yields is shown as box plots for all three RNA isolation protocols from PAXgene specimens combined (P, n = 45) and per protocol (PM, PQc, PQs, n = 15, each). The box plot shows the median values (center), the boxes represent the 25th and 75th percentiles, and whiskers indicate the 10th and 90th percentiles. Values outside the 10th and 90th percentiles are shown as individual data points. “*p*=*ns*” indicates differences between groups that are not significant with *p* ≥ 0.05.

The purity of RNA samples (UV260/280) was between 1.91 and 2.13 for EDTA samples and 1.98 and 2.10 for PAXgene samples. Purity was significantly higher for PAXgene compared to EDTA samples with the corresponding median purities of 2.02 and 1.97 (Fig 4).

**Fig 4.**
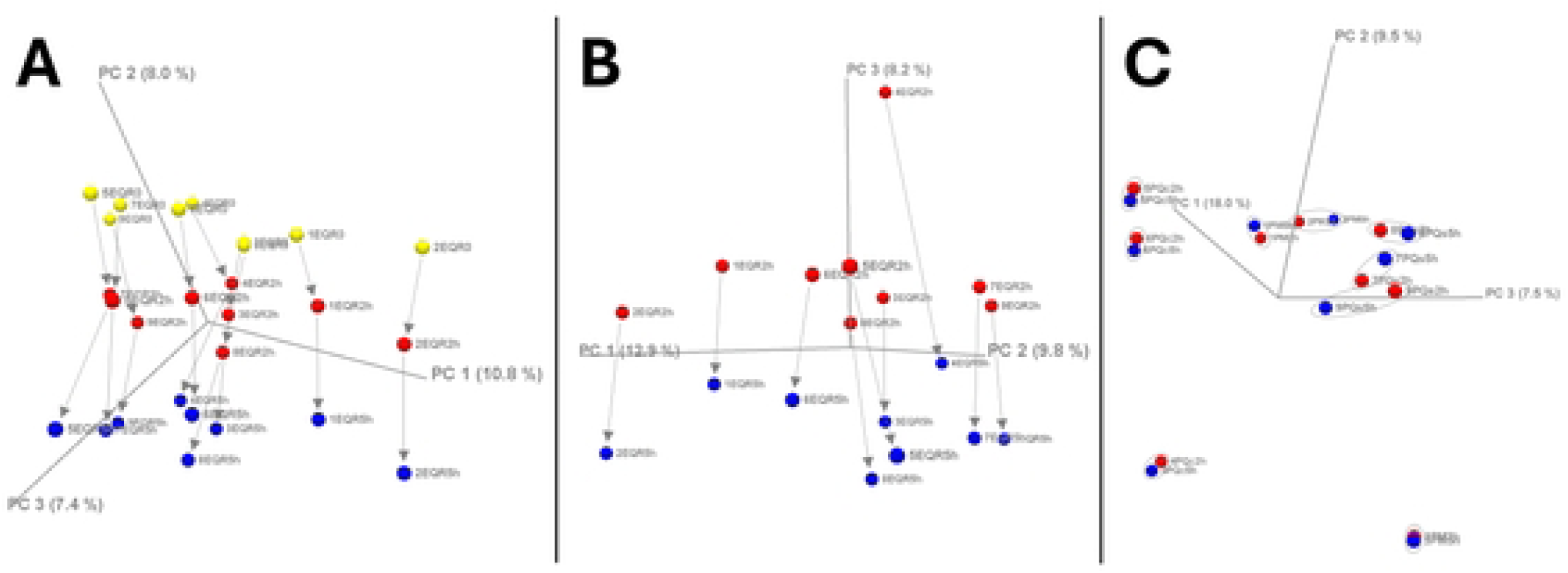
RNA quality control: RNA purity. RNA purity of all samples is shown as box plots with median values (center), 25th and 75th percentiles (box) and whiskers, representing the 5th and 95th percentiles. Values outside the 5th and 95th percentiles are shown as individual data points. E: EDTA tube specimens processed with EQR protocol (n = 54); P: PAXgene Blood RNA Tube specimens processed with PM, PQc and PQs protocol (n = 45). Statistical significance of difference is indicated as \*\*\*\**p* < 0.0001.

RNA integrity was statistically lower in EDTA than in PAXgene samples, with a median value of 8.5 (range: 7.4 to 9.0) compared to 8.8 (range: 6.7 to 9.9; Fig 5).

**Fig 5.**
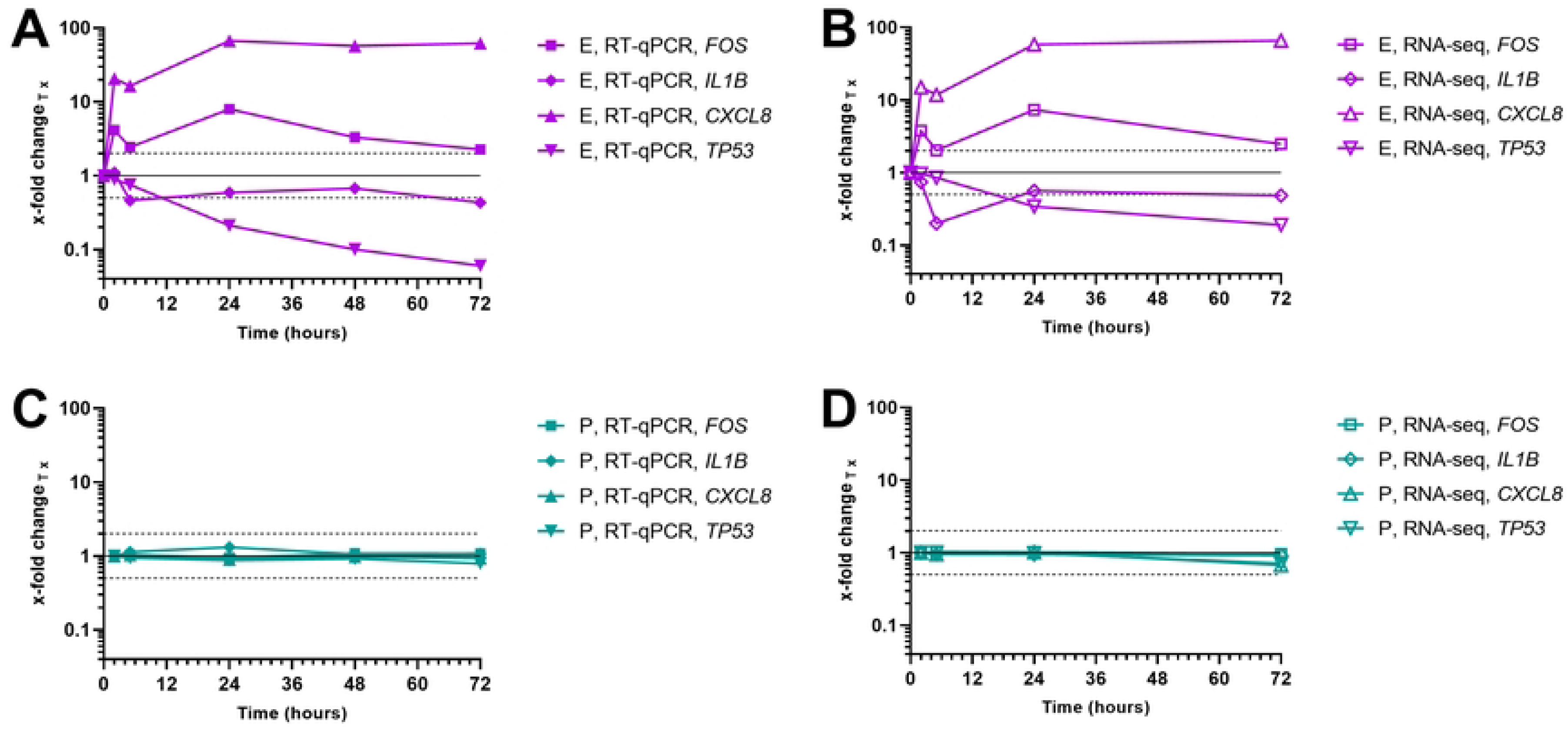
RNA quality control: RNA integrity. RNA integrity of all samples is shown as box plots with median values (center), 25th and 75th percentiles (box) and whiskers, representing the 5th and 95th percentiles. Values outside the 5th and 95th percentiles are shown as individual data points. E: EDTA tube specimens processed with EQR protocol (n = 54), P: PAXgene Blood RNA Tube specimens processed with PM, PQc and PQs protocol (n = 45). Statistical significance of difference is indicated as \*\**p* < 0.01.

For 87% of all PAXgene samples, the RNA integrity was higher than or equal to matching EDTA samples of the same subject with the same storage duration of blood specimen prior to RNA isolation (39 out of 45 sample pairs; Fig 6). In this direct comparison, PAXgene samples were on average 0.4 RIN higher than the corresponding EDTA samples.

**Fig 6.**
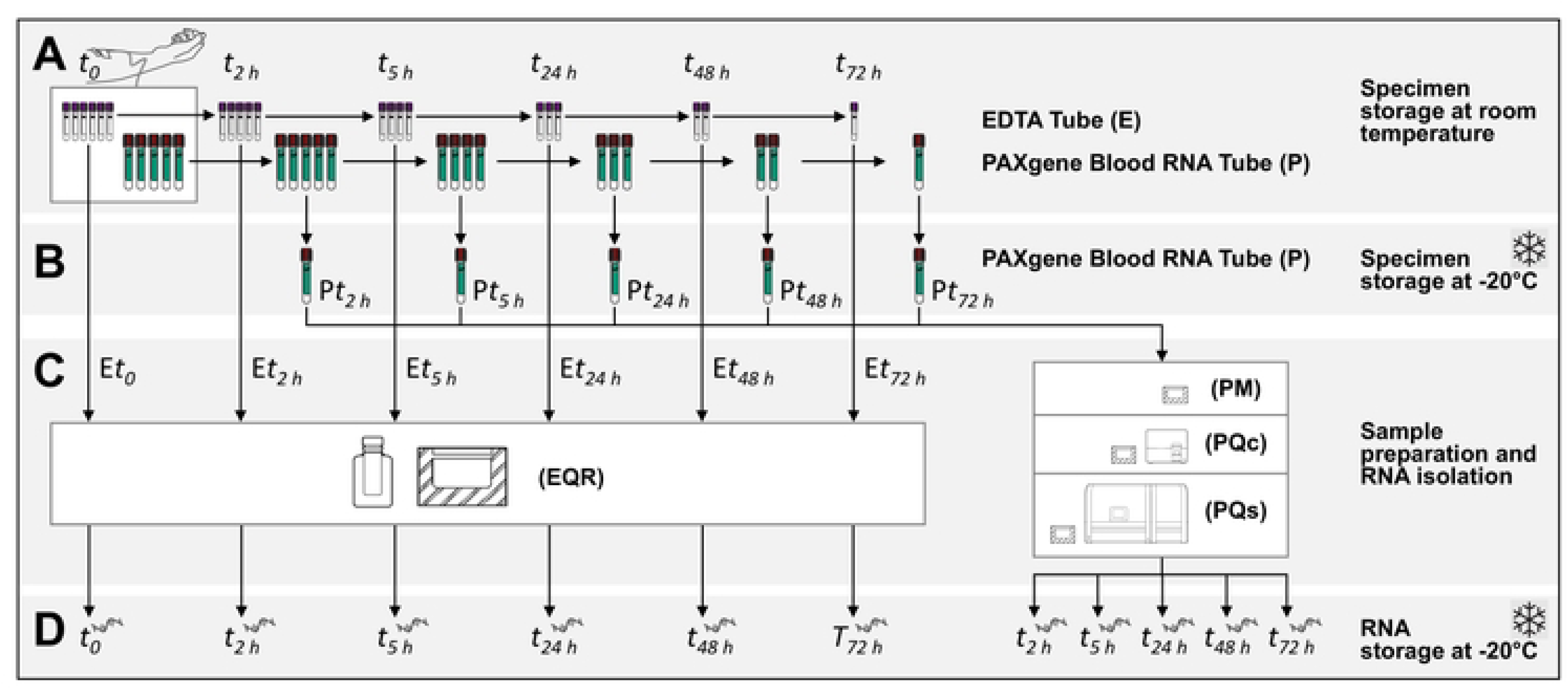
RNA quality control: RNA integrity comparison of matching samples. RNA integrity of EDTA and PAXgene sample pairs is shown, matching the same subject and storage period of blood specimens before RNA isolation. Individual data points are represented by dots, connected to corresponding data points of the data pair with lines (n = 45). Bars represent the means of both groups. E: EDTA tube specimens processed with EQR protocol, P: PAXgene Blood RNA Tube specimens processed with PM, PQc and PQs protocol (n = 45). Statistical significance of difference is indicated as \*\*\**p* < 0.001.

### Transcript level analysis by RT-qPCR

Transcript levels changed significantly during blood specimen storage in EDTA tubes for all genes analyzed by RT-qPCR; while for PAXgene blood, all changes stayed within the method’s precision for individual samples, available for *FOS* and *IL1B* (Fig 7A and B) and were near the baseline of zero for all transcripts (Fig 7A–D). Statistical tests confirmed that the differences measured for PAXgene samples were all statistically insignificant (*p* ≥ 0.05).

**Fig 7.**
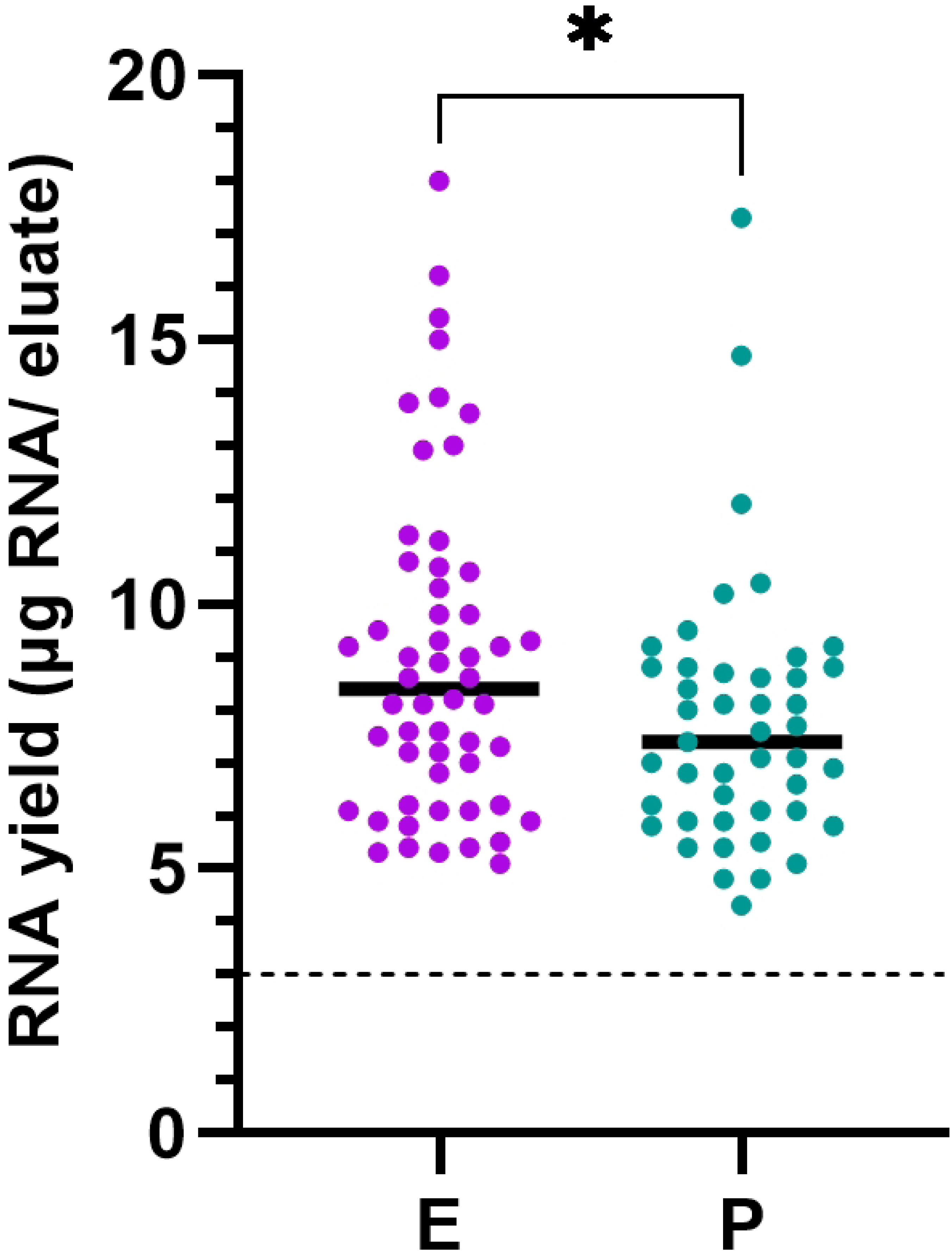
Transcript level differences as detected by RT-qPCR. Differences in gene transcript levels (A) *FOS*, (B) *IL1B*, (C) *CXCL8,* (D) *TP53* between reference samples (EDTA: E*t_0_*; PAXgene: P*t_2 h_*) and samples obtained from blood specimens incubated at room temperature for different periods of time up to 72 hours (3 days). Data are plotted as the mean ± SD of all subjects for EDTA (● purple circles, n = 9) and PAXgene (● turquoise circles, n = 9). Positive ΔΔCt (*FOS*, *IL1B*) and ΔCt values (*CXCL8*, *TP53*) indicate gains of transcripts, while losses are indicated by negative values. Dashed lines indicate the ± 3× total precision of the assays for single data points (A) ± 1.30 ΔΔCt, (B) ± 1.42 ΔΔCt. Levels of statistically significant differences are indicated as \**p* < 0.05, \*\**p* < 0.01, \*\*\**p* < 0.001 and \*\*\*\**p* < 0.0001 or are not shown if not significant (*p* ≥ 0.05).

Different courses and intensities of both increases and decreases of transcript levels were observed in blood specimens collected in the EDTA tubes of all subjects, depending on the analyzed target transcript and the duration of blood storage before RNA isolation (Fig 7).

No changes in gene expression levels were observed in PAXgene samples. In EDTA samples, transcript levels of two out of four genes of interest analyzed (*FOS*, *CXCL8*) were already significantly elevated after the shortest tested storage duration of 2 hours (Fig 7A, 7C). At this test timepoint, specimens from 89% of subjects (8 out of 9) showed *FOS* transcript levels increased above the method’s precision with a level of significance of *p* < 0.001 for the mean difference between reference (E*t_0_*) and test samples (E*t_2 h_*). At the same test timepoint, the difference for *CXCL8* was even greater and showed a higher level of significance, (*p* < 0.0001) with specimens of all subjects showing transcript levels increased above the method’s precision. Elevated transcript levels started to decrease slightly from 2 to 5 hours, only to rise again and reach a maximum after 1 day (24 hours), with samples from all subjects above the method’s precision for *FOS* and with a high level of significance of *p* < 0.0001 for the mean difference between reference (E*t_0_*) and test samples (E*t_24 h_*) for both *FOS* and *CXCL8*. The transcript level of *FOS* returned almost to baseline after 3 days (72 hours), while for *CXCL8* it remained consistently high and significantly increased (*p* < 0.0001).

The opposite trend was observed for *IL1B* and *TP53* (Fig 7B, 7D). Significant reduction of *IL1B* transcripts was first detected after 5 hours in 3 of 9 subjects affected (33%, *p* < 0.01 for the means), and for *TP53* after 1 day (24 hours, *p* < 0.01 for the means). For *TP53*, the losses continued steadily, reaching a minimum after the longest storage duration (72 hours, *p* < 0.0001), whereas for *IL1B*, the losses at 5 hours started to reduce until 24 hours, stayed at this level for a further day before dropping significantly to a minimum at the end of the time course (72 hours, *p* < 0.01).

*IL1B* and *CXCL8* showed the greatest differences in the magnitude of gene expression changes between subjects at individual test timepoints, which is reflected by the larger standard deviations compared to *FOS* and *TP53*. As an example, see *IL1B* E*t_48 h_*, E*t_72 h_* (Fig 7B) and *CXCL8* E*t_24 h_*, E*t_48 h_*, E*t_72 h_* (Fig 7C).

### Gene-expression analysis by RNA-seq

#### Assessment of rRNA and globin depletion

In blood samples, ribosomal RNA (rRNA) and globin mRNA represent the most abundant transcripts, with rRNA contributing up to 90% of total RNA and globin mRNA contributing up to 80% of protein-coding derived expression values [24]. Due to this ultra-high abundance, it is imperative to minimize the presence of rRNA and globin mRNA during stranded RNA library preparation to enable efficient usage of sequencing reads for robust expression analysis of protein coding genes. During NGS library preparation, QIAseq FastSelect -rRNA and QIAseq FastSelect -Globin were incorporated to deplete the presence of rRNA and globin mRNA. FastSelect blockers function by hybridizing tightly to the RNA target and preventing reverse transcriptases from synthesizing cDNA of the hybridized target. After sequencing analysis, the percentage of total reads that map to human rRNA and the percentage of FPKM that map to globin mRNA were measured. The results confirmed that FastSelect -rRNA and - Globin efficiently removed rRNA and globin respectively, with rRNA values ranging from 0.5% to a max of 3.7%, and globin mRNA values exhibiting a maximum of < 0.5% (data not shown). These low values ensured that the sequencing read budget was optimal for sensitive, robust whole transcriptome analysis, as the sequencing reads were not sequestered by the unwanted, high expression RNAs.

Analysis of NGS data using CLC Genomics Workbench revealed expression of approximately 18,000 genes per EDTA sample and 19,000 genes per PAXgene sample (FPKM > 0; data not shown). These values include both protein-coding and non-coding transcripts, indicating broad transcriptome coverage across both sample types and consistent library complexity.

### Gene expression concordance between different PAXgene RNA purification methods

Pearson correlation analysis was applied to assess whether different RNA purification methods introduced systematic changes in transcriptional profiles in the context of biological variability across healthy subjects [25]. Samples originated from different subjects and therefore represent independent biological observations rather than matched pairs. Correlations were calculated using RPKM expression values, with genes showing zero counts retained in the analysis to ensure unbiased comparison across the full expression range. Samples processed after 2 hours using either the QIAcube or QIAsymphony platforms showed high average Pearson correlation coefficients relative to manual preparation and to each other (0.98 ± 0.01, 0.97 ± 0.01 and 0.98 ± 0.01, respectively). These high correlations indicate strong overall similarity of gene expression patterns across subjects and demonstrate that transcriptional outcomes were consistent and comparable across purification methods, without evidence of systematic RNA isolation method-specific bias.

### Heatmap analysis of transcriptional stability

A heatmap of the most significant differentially expressed genes was generated using the top 1000 genes with a minimum of 100 counts and a significance threshold of p < 0.05 from blood samples collected from three independent subjects using either the EDTA or PAXgene workflow. This analysis identified 225 significantly deregulated genes in EDTA across subjects and incubation time points (Fig 8).

**Fig 8.**
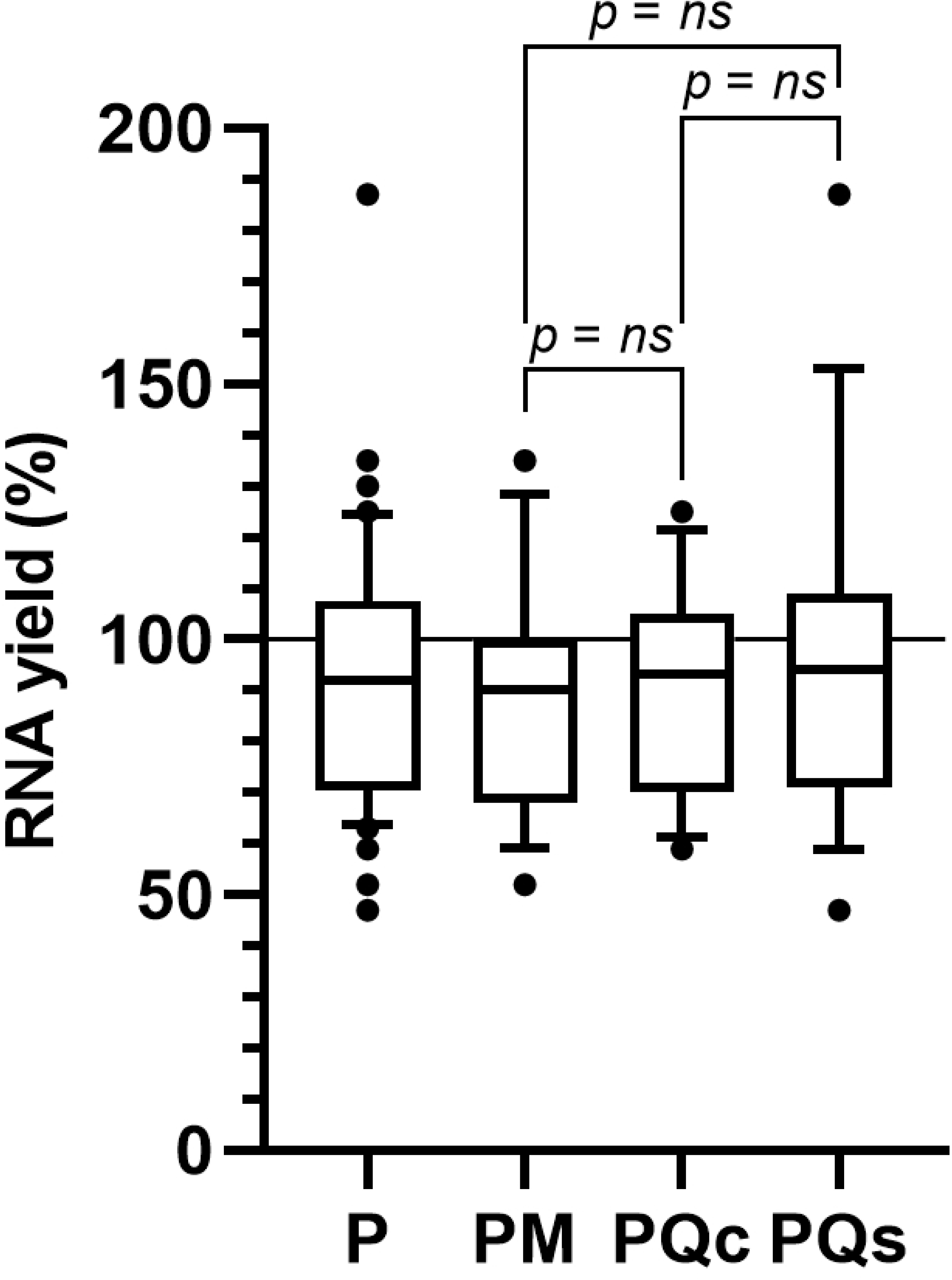
Heatmap of differentially expressed genes with hierarchical clustering. Heatmap showing the expression patterns of differentially expressed genes across the indicated subjects, workflows and blood storage time periods. Gene expression values were normalized and scaled by row (z-score). Rows represent samples and columns represent genes. Hierarchical clustering of genes and samples was performed using Euclidean distance and complete linkage, enabling grouping based on overall expression similarity and the formation of compact, well-separated clusters. The color scale indicates relative expression levels, with green representing higher expression and red representing lower expression. Only 225 genes meeting the significance criteria (adjusted *p* < 0.05 and | log₂ fold change | ≥ 1) are shown. On the left, EDTA samples are marked in purple and PAXgene samples in turquoise with slightly different color shades per subject. Sample names are coded by subject number (1, 2, 3), blood collection tube (E: EDTA, P: PAXgene), RNA isolation protocol (QR: QIAzol and RNeasy, M: PAXgene Blood RNA Kit, manual processing) and test timepoint given in hours (0, 2 h, 5 h, 24 h, 72 h).

Heatmap analysis revealed a clear separation between EDTA and PAXgene samples, as well as a pronounced time-dependent effect in EDTA samples. PAXgene samples clustered tightly across all subjects and storage time points, indicating highly stable transcriptional profiles with minimal variability over time. This clustering was consistent irrespective of subject identity, underscoring the effectiveness of PAXgene workflow in preserving RNA integrity and transcriptional states.

In contrast, EDTA samples exhibited progressive divergence from baseline expression profiles with increasing storage duration. While EDTA samples at early incubation times clustered relatively close to the reference condition (0 h, no storage), samples stored for 24 h and 72 h displayed pronounced shifts in gene expression and formed distinct clusters separated from both earlier EDTA time points and PAXgene samples. These patterns were consistently observed across all subjects, indicating that prolonged storage in EDTA tubes introduces systematic, time-dependent transcriptional alterations rather than subject-specific effects.

Overall, the heatmap analysis highlights the strong stabilization of transcriptional profiles achieved with PAXgene blood collection tubes and reveals a substantial, time-dependent transcriptional drift in EDTA samples, corroborating the results obtained from differential expression and volcano plot analyses (see Fig 9).

**Fig 9.**
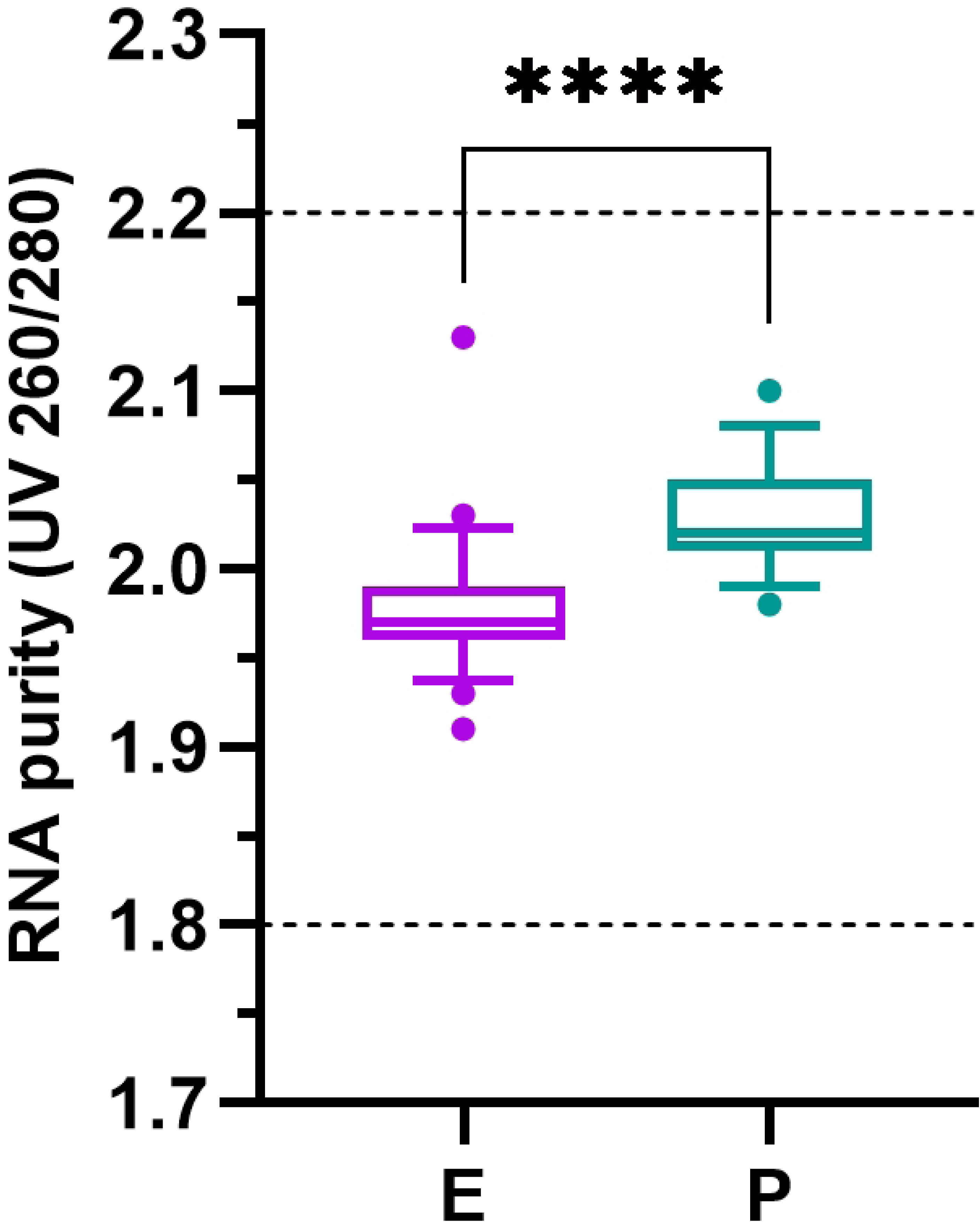
Volcano plot analysis of blood storage-induced gene expression changes in blood samples. Differential gene expression analysis was performed to assess the effect of blood sample incubation time while accounting for subject variability. Blood samples were incubated at room temperature for the indicated time points (2 h, 5 h, 24 h and 72 h) and analyzed in CLC Genomics Workbench using an ANOVA-like multi-group model in which incubation time was the primary factor. Subject effects were controlled by pairing samples from the same subject, thereby minimizing inter-subject variability and highlighting incubation-related expression changes. Tube-specific reference samples were used for comparison (EDTA: 0 h; PAXgene: 2 h). Significantly differentially expressed genes are shown as single dots for EDTA samples in (A) and for PAXgene samples in (B). Genes with increased expression are shown in green and genes with decreased expression in red, based on a fold-change threshold of ≥ 200% or ≤ 50% (| log₂ fold change | ≥ 1) and −log_10_ FDR p-value ≥ 1.3 (p < 0.05). Genes not meeting the significance criteria are shown in gray.

### Effect of blood storage time on transcriptional profiles

Volcano plot analysis revealed a strong time-dependent impact of blood storage at room temperature on gene expression profiles in EDTA samples (Fig 9A). At the earliest time point (2 h), only a limited number of genes were differentially expressed relative to the baseline reference samples (0 h, without storage), whereas prolonged storage resulted in a progressive increase in both the number and magnitude of deregulated transcripts. After 24 h and 72 h, EDTA samples exhibited widespread transcriptional changes, with a high number of genes showing significant upregulation (> 200%) or downregulation (< 50%), indicating substantial alterations to the transcriptomic profile over time.

Specifically, the number of deregulated protein-coding genes increased from 14 upregulated genes at 2 h to 42 upregulated and 14 downregulated genes at 5 h, followed by 488 upregulated and 350 downregulated genes at 24 h and reaching 971 upregulated and 1073 downregulated genes after 72 h, which represents 10.7% of the protein coding genes included in this analysis. In addition, deregulation of long intergenic non-coding RNAs (lincRNAs) was observed at 72 h incubation, with 45 deregulated lincRNAs out of a total of 2701 analyzed, representing 1.7% of the lincRNA population.

In contrast, PAXgene-stabilized samples demonstrated transcriptional stability (Fig 9B). Across all analyzed time points, including storage up to 72 h, no protein-coding genes exceeded the defined thresholds for differential expression, and no systematic shifts in gene expression were observed relative to the reference condition (2 h).

Together, these results demonstrate that room-temperature storage for different periods of time induces pronounced and cumulative transcriptional changes in EDTA-collected blood samples, whereas PAXgene tubes effectively preserve the transcriptional landscape over extended storage durations, supporting their use for robust and comparable gene expression analyses.

### Principal component analysis of tube type and incubation time

Principal component analysis (PCA) was performed to assess the impact of blood collection tube type and storage time on global gene expression patterns (Fig 10). At early blood incubation test time points up to 5 h, EDTA samples showed partial overlap. In contrast, at prolonged storage times (24 h and 72 h), EDTA samples formed distinct clusters according to incubation times that were clearly separated from earlier time points, whereas PAXgene samples of all blood incubation times clustered closely, including samples collected at earlier time points (0–5 h). All EDTA sample clusters formed stayed separated from the PAXgene sample cluster.

**Fig 10.**
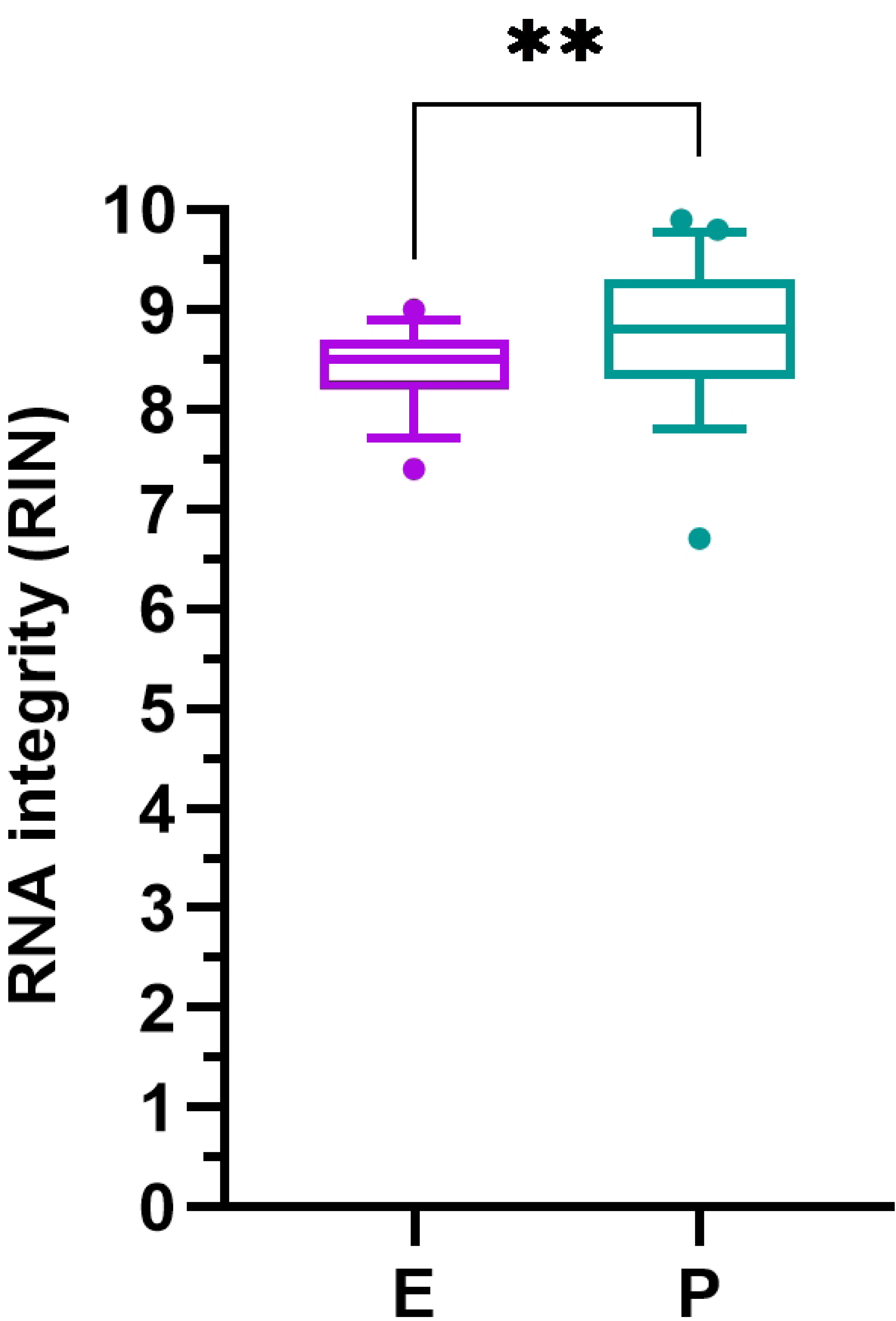
Three-dimensional PCA of gene expression changes during blood sample storage. 3D-PCA was performed on normalized gene expression data to visualize global transcriptional differences between blood samples stored for different periods of time at room temperature. The first three principal components (PC1, PC2 and PC3) explain the largest proportion of variance in the dataset. Each point represents an individual sample, with EDTA samples shown in purple and PAXgene samples shown in turquoise. Samples are plotted according to storage time points (0–72 h); the initial reference time point for PAXgene samples is 2 h due to processing requirements (see Materials and methods: time required for RNA complex formation and precipitation). After incubating blood for 5 h, EDTA samples of all subjects form separate clusters (24 h, 72 h), distinct from earlier time points as indicated by gray arrows, while PAXgene samples of all test time points cluster with samples from earlier storage time points of PAXgene. All EDTA samples cluster separately from PAXgene samples as indicated by a dashed gray line.

To further investigate patterns of global transcriptional differences occurring within the first 5 hours post-collection, 3D-PCA was performed with a subset of data of all nine subjects (Fig 11). In samples collected in EDTA tubes, a subject-independent pronounced and consistent shift in gene expression pattern from collection timepoint (0 h) samples across the shortest test time point analyzed (2 h) was observed that continued to the 5 h time point (Fig 11A). To enable direct comparison with PAXgene samples, the 0 h EDTA time point was further excluded; nevertheless, a comparable shift in gene expression was observed as incubation time increased from 2 h to 5 h (Fig 11B). In contrast, PAXgene samples did not exhibit a shift in gene expression patterns from 2 h to 5 h incubations, indicating stable transcriptional patterns over this time interval (Fig 11C).

**Fig 11.**
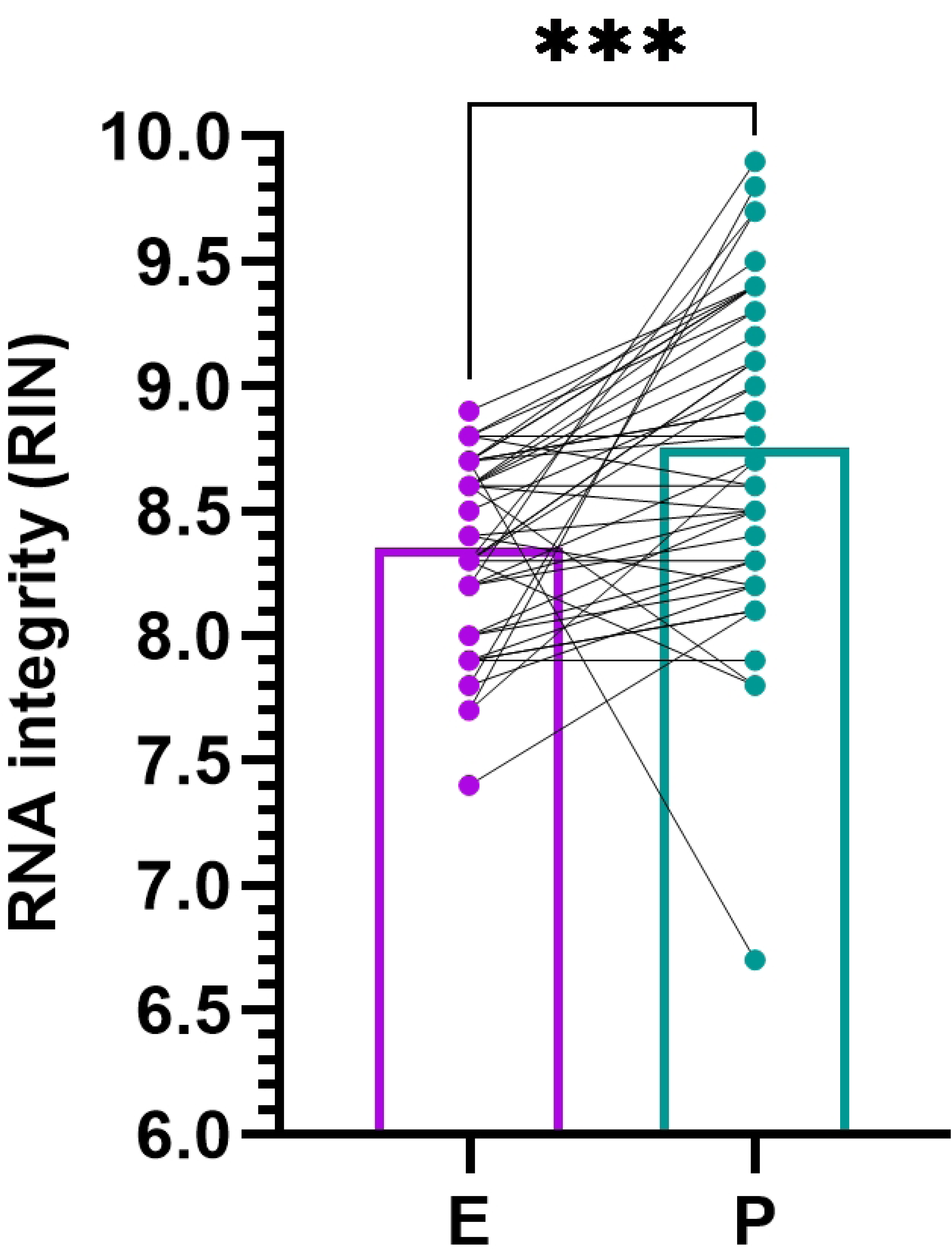
Three-dimensional PCA of gene expression changes during the first 5 hours post-collection. 3D-PCA was performed as described in Fig 10 on normalized gene expression data to visualize global transcriptional differences in EDTA (A, B) and PAXgene samples (C) stored for up to five hours at room temperature after blood collection. Samples were analyzed at the indicated time points, shown in yellow (t0, EDTA), red (t2 h, EDTA and PAXgene) and blue (t5 h, EDTA and PAXgene). In EDTA sample sets, arrows indicate a consistent shift in gene expression patterns from the initial time point (t0, no incubation) across 2 h and 5 h. In PAXgene sample sets, ovals indicate that paired samples that were incubated for 5 h cluster very closely with the 2 h reference samples from the same subject.

Taken together, these analyses demonstrate that transcriptional variability in EDTA samples is primarily driven by storage time, whereas PAXgene tubes preserve stable and comparable gene expression profiles across sample incubation periods.

### Comparison of gene expression analysis between RT-qPCR and RNA-seq

The direct comparison of transcript level changes as the effect of blood sample storage revealed a high degree of agreement between RT-qPCR and RNA-seq with similar time courses of increased and decreased transcript amounts with EDTA samples and no changes in PAXgene samples (Fig 12).

**Fig 12:**
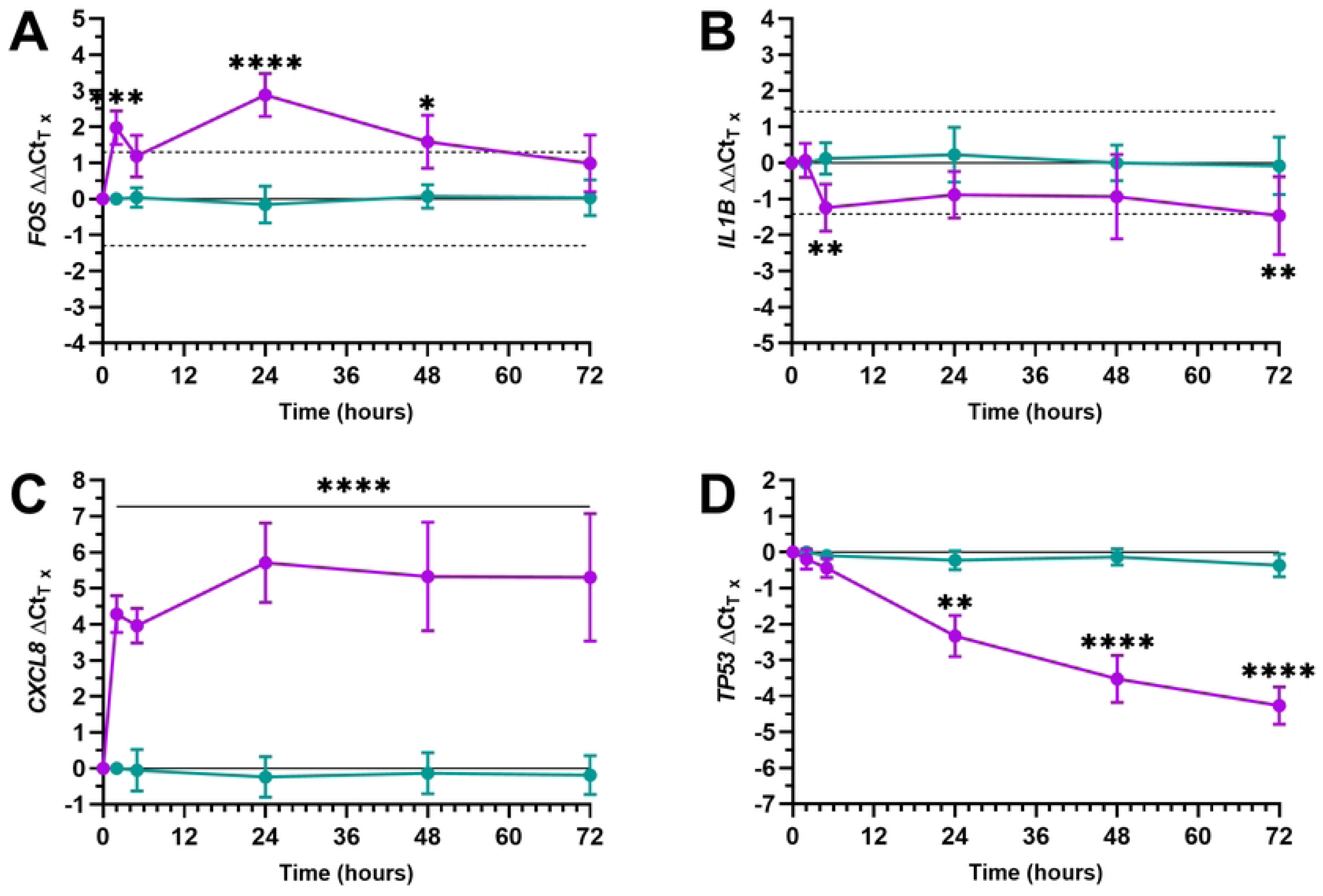
Comparison of transcript level changes quantified by RT-qPCR and RNA-seq. *FOS, IL1B, CXCL8 and TP53* transcript level changes in blood specimens as the result of blood storage at room temperature were calculated as multiplication by x (x-fold changes) based on (A, C) RT-qPCR (ΔCt, ΔΔCt; filled symbols) and (B, D) RNA-seq data (number of reads per target; open symbols) and are plotted as means from all subjects (n = 9). Specimens were collected in (A, B) EDTA (purple) and (C, D) PAXgene Blood RNA Tubes (turquoise) and incubated for the indicated periods of time up to 72 hours (3 days) before RNA isolation. The baseline of 1 indicates no change, while gains and losses of transcripts are indicated by values > 1 and < 1, whereby double (2-fold-change) and half of the initial transcript amount (0.5-fold-change) are highlighted by dashed lines.

After 2 hours, the *CXCL8* level had already increased by a factor of 21 (RT-qPCR) or 15 (RNA-seq), and that of *FOS* by a factor of 4 (RT-qPCR and RNA-seq). After 5 hours, the *IL1B* level was only 20% and 46% (0.2- and 0.46-fold change, RNA-seq and RT-qPCR, respectively) of the initial amount. The highest increase was measured for *CXCL8*, with a 67-fold change after 1 day (RT-qPCR) and a 66-fold change after 3 days (RNA-seq), and for *FOS* after one day with an 8-fold (RT-qPCR) and 7-fold change (RNA-seq). The greatest loss of transcripts was observed for *TP53*, with a reduction to 7% and 19% of the initial level after 3 days (0.07 and 0.19-fold-change, RT-qPCR and RNA-seq, respectively).

## Discussion

### Overall RNA yield and purity

RNA yields were comparable between EDTA and PAXgene tubes within overlapping ranges, with slightly higher median yield in EDTA. This aligns with prior work by Rainen et al. [26] showing that PAXgene stabilizes RNA without substantially compromising yield, despite differences in extraction chemistries.

RNA purity was higher in PAXgene samples compared to EDTA samples, independent from different RNA isolation chemistry and protocol applied to PAXgene samples (silica membrane, manual and automated and automated magnetic bead-based methods). The direct comparison of EDTA and PAXgene samples isolated with the same principle of RNA isolation chemistry after cell lysis, using chaotropic salts, ethanol, binding to silica membrane, enzymatic on-column DNA digestion, washing off impurities and elution of the RNA from membrane in a spin column format showed higher purities for PAXgene. Malentacchi et al. [27] showed superior purity and reduced genomic DNA contamination in stabilized PAXgene samples, while EDTA samples showed more handling-induced variability.

#### RNA integrity (RIN)

Higher RIN values in PAXgene samples indicated better preservation of RNA during pre-analytical storage and processing. PAXgene tubes improved RNA integrity when comparing EDTA and PAXgene samples matched for blood storage times. The improvement is consistent with Martire et al. [4] who reported that stabilization chemistry prevented rapid degradation seen in EDTA after short bench times of 2 h. Tang et al. [28] reported long-term evidence of excellent RNA stability in PAXgene tubes over 7–11 years of cryostorage, supporting robustness for NGS workflows.

#### Impact of rRNA and globin depletion during RNA-seq library preparation on transcriptome coverage

Efficient depletion of rRNA and globin transcripts has proved to be essential for maximizing sequencing efficiency and transcript recovery [29]. The low residual rRNA and globin mRNA levels observed here indicate effective suppression of ultra-abundant transcripts, enabling deep and uniform coverage of the transcriptome. Detection of approximately 18,000–19,000 expressed genes per sample confirms that the FastSelect probe-based depletion supports sensitive whole-transcriptome analysis without apparent loss of complexity or systematic bias [30].

### Concordance of gene expression across RNA purification workflows

RNA purification with PAXgene samples, including automated sample processing workflows, had only a negligible impact on gene expression measurements. High Pearson correlation of RNA-seq data across different RNA isolation methods for PAXgene samples, even when genes with zero counts were included, indicated that transcriptional patterns were not dominated by method-specific effects. This observation is in line with large-scale RNA-seq benchmarking studies showing that, under controlled conditions, different RNA extraction and processing workflows yield highly concordant global expression profiles and that Pearson correlation is a sensitive metric for detecting technical bias in replicate comparisons [25,31]. Taken together, the high concordance observed across RNA isolation workflows for PAXgene indicated that these are suitably harmonized and technically robust, making downstream biological interpretation unlikely to be affected by the choice of isolation method used in this study.

### Pre-analytical stress: Blood storage duration has an impact on gene expression in EDTA, not in PAXgene samples

Drastic ex vivo changes in transcription were found and confirmed in EDTA blood during 2–72 h of storage at room temperature by RT-qPCR and RNA-seq, with *FOS* and *CXCL8* transcripts increased sharply up to 7–8 and 66–67-fold and *IL1B* and *TP53* significantly decreased down to a minimum of 2.2–5.0 and 5.3–14.3-fold over time.

Blood samples stored in PAXgene tubes demonstrated no significant changes over blood sample storage, with all differences staying within RT-qPCR assay precision and below FDR applied in RNA-seq – confirming the successful suppression of ex vivo transcriptomic changes that was also reported by Rainen et al. [26]. Pahl & Brune [32] reported activation of leukocytes in EDTA blood samples during pre-analytical delays with *IL1B*, *IL6* and *TNF* transcripts rising over hours after phlebotomy.

Bench-time is a dominant confounder as strongly supported by multiple studies showing delayed EDTA processing affects transcriptomes even at 4°C [4,33] and shipping and transport conditions alter expression patterns in EDTA blood, whereas PAXgene blood showed minimal effects [34]. Wang C. and Liu H. [35] found that storage duration of fresh whole blood at room temperature had the largest effect on RNA degradation that was higher than the effects of cDNA storage at −20°C and storage of isolated RNA at room temperature with an average half-life of mRNA analyzed of 16.4 h. Johnson et al. [36] found by immunophenotyping that cellular profiles of unstabilized blood stored and shipped at ambient temperature for 24 h were preserved, although some populations were broadened and CD16 intensity was decreased on classical monocytes. With RNA-seq analysis they identified that a transcription factor module associated with inflammation was upregulated. Cooling to 4°C and shortening the time of sample storage at RT to few hours does not change the expression of selected genes in unstabilized blood as reported by Schüle et al. [37].

In contrast, Xing et al. [38] reported that storage of EDTA samples for 24 h at 4°C and room temperature affected transcript levels of 1515 (1.51%) and 10,823 (10.82%) genes in isolated leukocytes, respectively, when compared to fresh leukocytes. Upregulated genes were involved in apoptotic signaling pathway, negative regulation of cell cycle and lymphocyte activation.

Zhang et al. [5] leveraged ex vivo-induced changes in EDTA blood to identify and validate, by means of microarray analyses, transcripts that are suitable as marker transcripts for use in quality control assays to qualify RNA for other downstream assays.

### Effects of blood collection tube type and blood storage dominates over subject

The choice of blood collection tube and blood storage duration ahead of RNA isolation emerged as the dominant sources of transcriptomic variability. EDTA samples exhibited pronounced, time-dependent transcriptional drifts that affected a substantial fraction of the transcriptome. The consistency of these alterations across subjects suggests that storage duration is the primary contributing factor, rather than the inter-individual variability (Fig 8).

Previous RNA-seq–based studies of EDTA-collected blood have similarly reported that even short pre-analytical delays result in widespread differential expression, particularly affecting inflammation- and stress-responsive transcripts [4,39]. The involvement of both protein-coding and long non-coding RNAs observed here indicates a broad disruption of transcriptional homeostasis, as has been noted in other investigations of suboptimal blood handling conditions [40].

3D-PCA revealed a consistent, time-dependent transcriptional shift in EDTA samples, whereas PAXgene samples showed stable expression profiles under all conditions. In EDTA samples, gene transcripts behaved very differently across the different time points (increasing/decreasing transcript levels, induction and loss, etc.). Such shifts were not observed in PAXgene samples. A previous study of blood transcriptomes that used microarray analysis has reported comparable PCA-based differences between EDTA and PAXgene blood, mainly caused by pre-analytical delays and leukocyte isolation from EDTA [41]. The tight clustering of PAXgene samples observed in the present study confirms that immediate RNA stabilization effectively arrests transcriptional activity and preserves biological states, consistent with extensive prior evidence also demonstrating long-term transcriptome stability in PAXgene tube-stabilized whole blood stored frozen for months and years [26,28,41].

#### Comparison of RT-PCR and RNA-seq

The study demonstrates high concordance between RT-qPCR–based ΔCt and ΔΔCt changes and RNA-seq fold-changes across genes tested. This supports the analytical validity of RNA-seq for detecting early transcriptomic instability when pre-analytical processing is suboptimal. Concordance aligns with prior findings from Donohue et al. [42] who analyzed the impact of blood collection tubes on expression signatures, showing RNA-seq robustness provided RNA integrity is maintained, reinforcing the requirement for using RNA stabilization tubes.

#### Biological interpretation and Implications for research, development and diagnostics

The strong transcriptional instability observed in EDTA samples is biologically plausible. In the absence of immediate RNA stabilization, blood cells remain metabolically active outside the subject’s body.

Unlike PAXgene tubes that arrest transcriptional activity and stabilize RNA at the time of blood draw by immediate cell lysis causing cell death, cells in EDTA tubes – particularly leukocytes – remain metabolically active for several hours ex vivo. During this period, cells may respond to multiple stressors during sample storage and handling. Calcium chelation induced by EDTA, mechanical stress, temperature changes, hypoxic conditions, nutrient deprivation and accumulation of metabolites and other factors are likely to trigger stress-responsive and inflammatory pathways and initiate early apoptotic programs. These ongoing cellular responses likely contribute to the progressive transcriptional changes observed with prolonged storage in EDTA tubes [4].

Regulation of genes such as *FOS*, *CXCL8*, *IL1B* and *TP53* is consistent with leukocyte activation and stress signaling, which have been shown to arise rapidly after blood collection in EDTA tubes and to intensify with prolonged handling times [4,26,32]. In contrast to the viable cells remaining in EDTA tubes, cells in PAXgene Blood RNA Tubes are lysed by the cationic detergent contained in the RNA stabilization solution, thereby effectively inhibiting gene regulation. In addition, the detergent binds to RNA and thus prevents enzymatic degradation by RNases present in blood [43], and precipitates the RNA, enabling its subsequent isolation from a reduced sample volume following blood tube centrifugation.

Pre-analytical changes observed in EDTA blood samples can obscure genuine biological signals, thereby hindering research into gene expression regulation, alterations and differences in various contexts, as well as the discovery, verification and validation of blood RNA-based biomarkers. In the worst case, these changes could lead to false positive or negative results in blood RNA-based diagnostics.

For reliable gene expression analysis of blood RNA, standardizing workflows using immediate RNA stabilization during blood collection and quality control is key, as shown here and emphasized in the international guideline ISO 20186-1 [21] for the pre-analytical phase of RNA blood testing. Proficiency testing PT (EQA) may serve as a tool to assess the level of control achieved over pre-analytical variables of blood RNA workflows [20,27,44].

### Strengths of this study

Extensive transcriptional changes were identified in unstabilized whole blood samples without isolation of leukocytes prior to RNA isolation (bulk RNA), compared with stabilized samples from the same subjects under identical sample storage conditions using a time-course experiment. Relative gene expression changes measured by RT-qPCR were confirmed by RNA-seq analysis. All differentially expressed protein-coding genes and the magnitude of changes were identified.

The effects caused by blood storage durations as they occur in real-world settings, including blood RNA biomarker discovery studies, clinical trials and diagnostic workflows were investigated. This included both shorter blood storage intervals (2 h, 5 h), for example when blood collection and RNA isolation are performed within the same hospital, or when blood is collected from multiple patients in the morning, samples are pooled until noon, and RNA is isolated in batches in the afternoon; as well as longer storage and transport times (1 to 3 days), when sample collection and processing are geographically separated and samples require shipment, such as in clinical multicenter studies or diagnostic testing performed in distant reference or central laboratories. This is especially critical if the test group and control group are collected at different sites with different pre-analytical sample logistics, e.g., different specimen storage and transport times.

### Limitations

The gene panel analyzed by RT-qPCR (*FOS, IL1B, CXCL8, TP53*) captured only limited pathways. This initial limitation was overcome by applying RNA-seq on the same samples, enabling a broader transcriptome-wide view and confirming the observed effects. The storage conditions modeled room temperature but no other scenarios (e.g., shipping at 4°C, fluctuating temperatures, elevated temperatures above RT).

Since changes in EDTA blood were already detected at the first testing time point (2 h), it remains unclear when the earliest measurable alterations actually occur.

In general, bulk RNA analysis as reported here may obscure cell-type dynamics caused by a specific type of blood cells that are too small to be measured early in the background RNA from cells not in focus. This can be resolved by analyzing the transcriptomes of individual cells. Considering that the pre-analytical phase of EDTA samples causes ex vivo gene expression changes as reported and additional changes are induced by the isolation of PBMCs [45], stabilization is also key for transcriptomics on a single cell level. Single-cell analysis is not possible with PAXgene samples as the RNA stabilization reagent in the tube lyses the cells, but cell nuclei from PAXgene tubes remain intact enabling single-nuclei RNA-seq with comparable gene expression profiles to PBMCs, isolated from matching samples [46].

## Conclusion

Our study fills a critical gap by providing the first systematic RNA-seq analysis of how gene expression profiles in human EDTA blood stored at room temperature change and diverge over time. Despite the continued use of EDTA blood in numerous gene expression studies, there has been a lack of quantitative data characterizing the transcriptome-wide extent, kinetics and biological implications of ex vivo transcriptional drift in these samples. By addressing this, our work is essential for understanding and defining the dynamic nature of gene expression changes in unstabilized blood, providing valuable insights for both research and clinical applications.

Our results clearly demonstrate that EDTA tubes are unsuitable for transcriptomics studies with multi-hour pre-analytical delay due to rapid, dramatic transcriptomic drift.

In contrast, PAXgene Blood RNA Tubes in combination with different PAXgene Blood RNA isolation kits providing manual and automated sample processing options and optimized for the tube maintain RNA quantity, purity, integrity and transcriptome stability, supporting their use as the pre-analytic gold standard for blood-based molecular RNA assays using RT-qPCR and RNA-seq.

The findings align with and reinforce more than two decades of evidence supporting RNA stabilization systems as essential for reproducible human transcriptomics, by applying state-of-the-art technology RNA-seq.

## Acknowledgements

We thank all donors who provided blood specimens for this study, Franziska Heese and Maike Schönborn for excellent technical assistance, Markus Wolf for the support in the selection and application of statistical tests on RNA quality, quantity and RT-qPCR data and

Emma Waldron Smythe, who provided medical writing services on behalf of QIAGEN GmbH. The authors are grateful to the EU FP7 project SPIDIA (grant no. 222916) and the EU H2020 project SPIDIA4P (grant no. 733112) for the efforts made to create ISO standard 20186-1 for the pre-analytical phase of blood RNA that was applied in the study.

